# Gene editing without a genome: generation and validation of F0 CRISPR mutants in gastropod mollusc *Crepidula fornicata*

**DOI:** 10.1101/2025.11.26.690768

**Authors:** Danielle C. Jordan, Molly L. Rivers, Lucy Belmont, Samuel A.M. Martin, Tim P. Bean, Victoria A. Sleight

## Abstract

**Background:** CRISPR-Cas9 gene editing is a powerful tool to study gene function but to date is mostly applied in traditional model organisms. Due to specific challenges in spiralian genomics such as genome size, complexity, proportion of repetitive regions and high inter individual genome variation, applications of CRISPR-Cas9 in spiralian models have been limited. Within the Spiralia, molluscs are a strikingly diverse phylum with many unique gene family expansions and novelties, yet there are relatively few applications of CRISPR-Cas9 to unravel gene function.

**Results:** We generated *pax6* knockout F0 CRISPR-Cas9 mutants in the gastropod mollusc *Crepidula fornicata* using a *de novo* transcriptome to design single guide RNAs and genotyping primers. In lieu of an assembled genome, alignments with closely related species were used to determine putative intron-exon boundaries and successfully target gene editing to a specific exon of *pax6* containing a homeodomain. F0 *pax6* knockout mutants had an eye-loss phenotype. Successful CRISPR-Cas9 gene editing was confirmed genotypically using Sanger sequencing and Interference of CRISPR Edits (ICE) analysis.

**Conclusions:** The generation of F0 CRISPR-Cas9 knockout mutants with a clear phenotype in a mollusc without an assembled genome enhances the applicability of CRISPR for functional genomics in non-traditional emerging model systems. *pax6* is highly conserved and is required for eye development across metazoans, including the snail *C. fornicata*. The *pax6* mutants developed in this study suggests the *pax6* homeodomain is specifically required for eye formation in gastropods and sheds light on the evolution of eye development in animals.

## Background

For more than a decade, CRISPR-Cas9 gene editing has been used as a tool to study gene function in traditional model organisms such as fly, mouse, and zebrafish (1). To date however, CRISPR-Cas9 gene editing has not been routinely applied in spiralians. Spiralia are a morphologically and ecologically diverse group comprising nearly half of extant metazoan phyla and around 10% of known animal species. There are a range of reasons it is challenging to conduct CRISPR-Cas9 experiments in spiralians. Firstly, species are rarely routinely kept in the lab and so animal husbandry protocols require extensive optimisation to access one-celled zygotes. Delivery of CRISPR-Cas9 reagents to embryos via microinjection or electroporation is also challenging due to encapsulation or embryo viability after such manipulations. Viral delivery methods, such as lentivirus that can be modified to infect invertebrate cells, have not been established for spiralians (2). In many spiralian groups, high quality chromosome-resolved genomes have been challenging to assemble due to the unique genome biology of some taxa, such as molluscs. Molluscs produce a polysaccharide-rich mucus that make it difficult to obtain high molecular weight DNA needed for the long-read sequencing required for genome assemblies (3). Beyond obtaining high-quality DNA, mollusc genomes have highly repetitive sequences and high rates of heterozygosity that short reads alone cannot resolve (4,5). As the number of available spiralian genomes are increasing, largely due to access to long-read technologies, CRISPR-Cas9 gene editing is beginning to be utilized.

Since 2020, there has been an uptick in the successful usage of gene editing in spiralians. In the platyhelminth *Schistosoma mansonii*, electroporation was used to transfect one-celled zygotes with a CRIPSR vector using homology directed repair (HDR) to conduct both KO and knock-in experiments (6). In the marine annelid *Capitella teleta* CRISPR-Cas9, delivered via microinjection, was used to conduct KO experiments (7). In the Farrer’s scallop, *Chlamys farreri,* and the Yesso scallop, *Mizuhopecten yessoensis,* gene function has been studied *in vitro* via primary cell lines with electroporation as a delivery mechanism (8,9). A new technique called VitelloTag has been developed and applied in deuterostomes (the bat star, *Patiria miniata* and the acorn worm, *Saccoglossus kowalevskii*) and promises to be applicable to any metazoan excluding ctenophores (10). VitelloTag utilizes the conserved yolk protein, vitellogenin, by synthesizing a receptor-targeting peptide fused with a single-guide RNA (sgRNA) and Cas9 protein that can be incubated with oocytes and taken up through receptor-mediated endocytosis (10). This new method bypasses the need for microinjection and electroporation, though is currently limited to metazoans where *in vitro* fertilization is possible, it has not yet been demonstrated in spiralian taxa or achieved complete KO efficiency (10).

Possibly the most iconic of the spiralian clades, and certainly the most speciose, are the molluscs. Molluscs are the second most speciose animal phylum, and due to their huge diversity, no one species is accepted as a representative model system (11). Most CRISPR-Cas9 studies in molluscs build on foundational work and microinjection techniques developed in the slipper limpet, *Crepidula fornicata* where the first CRISPR-Cas9 mediated genome editing in a mollusc was achieved via knock-in (12). HDR was used to fuse mCherry fluorescent protein to β-catenin and successful CRISPR-Cas9-mediated gene editing was validated phenotypically through visualisation of mCherry; however, genotyping at the gDNA level wasn’t performed, and authors inferred successful genome integration of the knock-in at the RNA level (12). Since 2014, there are several examples of F0 crispant KOs in molluscs, all using microinjection as the delivery system: *tryptophan 2,3 dioxygenase* in *Doryteuthis pealeii* (13), two *tyrosinase* genes in *Magallana (*formerly *Crassostrea) gigas* (14), *calaxin* in *Lottia goshimai* via microinjection (15), and most recently, *CaSMP1* in *Crepidula atrasolea* (16). There are also two examples of mollusc F2 null KOs, for *pax6 in Pomacea canaliculata* (17) and *lsdia1* in *Lymnaea stagnalis* (18).

*C. fornicata* has served as a spiralian embryological model for 150 years (19–21). *C. fornicata*’s developmental model status is due to its ability to be easily cultured in lab, large number of embryos (22), and amenability to embryonic manipulations such as microinjection (12), ablation (23) and live imaging (24). High-resolution lineage tracing in *C. fornicata* has produced a detailed fate map for this species and the first spiralian CRISPR-Cas9 gene editing example (12,19,24–27). Despite its contributions and prominence as an embryological model system, *C. fornicata* does not have an assembled genome. To date, a lack a genome has limited *C. fornicata*’s use to answer genetic questions. Recent research efforts have moved towards a close relative, *C. atrasolea*, due to ease of laboratory culture owing to its direct development and short generation time compared to *C. fornicata,* and assembled genome that is currently not publicly available but referred to in a recent publication (16,21,23). *C. fornicata* remains a tractable model for genetic manipulation due to embryonic robustness to microinjection and large brood size compared to *C. atrasolea* (12,22). Significantly, having multiple genetically enabled, closely related species within a genus that captures interesting developmental and ecological diversity, provides a powerful group for eco-evo-devo studies in molluscs.

Positive control genes are necessary to assess the efficacy of CRISPR-Cas9 and form an essential step in method optimisation. Positive control genes should provide a consistent, easy-to-interpret phenotype (28). Pigmentation genes, like tyrosinase (*Tyr*), are often chosen as positive control for KO experiments, for example *Tyr* null mutants in mouse produces albinism (29,30). While present in molluscs, *tyrosinase* is thought to be pleiotropic with roles in both pigmentation and biomineralization limiting its use as a solely pigmentation-related positive control gene target (14,31). In the cephalopod *Doryteuthis pealeii*, knocking out *tryptophan 2,3 dioxygenase* resulted in reduced pigmentation in retinas and chromatophores (13), demonstrating that albino phenotypes as positive controls and proof-of-concept KOs are achievable in molluscs. Here, we chose the eye development gene *pax6* as a positive control for CRISPR-Cas9 KO proof-of-concept in *C. fornicata*. While likely possible to achieve, a pigmentation gene KO was not chosen due to the diversity of pigmentation mechanisms in molluscs (32). Instead, we rationalised that the transcription factor *pax6* may have a more conserved function across metazoans and be more straightforward to phenotype. Present in all genomes of animals with a visual system, *pax6* is considered a primary regulator of eye development (33). KOs of *pax6* disrupt eye development and cause an eyeless, or reduced eye, phenotype across many invertebrate and vertebrate model organisms (34,35). In the gastropod, *Pomacea canaliculata*, a deletion in the Pax domain from the *pax6* gene produced an eyeless phenotype in F2 homozygous mutants (but no phenotype was seen in any F1s or F0s) (17). Because the *pax6* KO eyeless phenotype is well described and consistent across metazoans, it can serve as an effective control gene to establish the efficiency of CRISPR-Cas9 KO in *C. fornicata* to develop a feasible and reproducible KO experimental pipeline. In addition, in *C. fornicata*, the larval eyes develop by the late organogenesis stage in the form of two dark spots seen from an anterior view (19), meaning KO mutants can be screened from an early stage.

From simple pit eyes to complex camera-type eyes, mollusc visual systems are the most diverse in form of any animal phyla (36), and represent a powerful system to understand the repeated evolution of complex organs (37). An elevation of tools and expertise in molluscan model systems however, is required to tackle the genetic, cellular and developmental control of eye formation in multiple comparative species (38). Most recently, *P. canaliculata* has been proposed as a genetically tractable molluscan system to study complete camera-type eye regeneration (17). *P. canaliculata* has impressive eye regeneration capabilities and its embryos can now be collected, microinjected with exogenous mRNA or CRISPR-Cas9 reagents and cultured *ex ovo.* While *C. fornicata,* to date, has not received research attention on its visual systems, its history as an embryological model could make it a compelling comparative caenogastropod species for molluscan eye evo-devo studies.

In this paper, we sought to demonstrate the feasibility of gene-editing in molluscs explicitly using a *de novo* transcriptome to design gRNAs and genotype. Compared to a genome, assembling and annotating a transcriptome is less expensive and technically challenging (39), making transcriptomes more feasible and accessible to a wide range non-traditional model systems. In a phylum with over 100,000 described extant species, there are now approximately 1,000 publicly available mollusc genome assemblies, including around 250 to 300 at chromosome-level resolution and roughly 500 at scaffold or contig level in NCBI GenBank as of 2025 (40). There are also approximately 850 to 1,100 assembled molluscan transcriptomes publicly available in the NCBI Transcriptome Shotgun Assembly (TSA) database, along with around 40,000 raw RNA-seq datasets in the Sequence Read Archive (SRA) that could be used to generate additional transcriptome assemblies (41,42).

Not just an advantage for feasibility, using a transcriptome to design gRNAs has several benefits. Molluscs are genetically diverse at the population and inter-individual level (43–46). For example, there is significant and persistent genetic diversity in the highly dispersive blue mussel genus, *Mytilus*, where species hybridization is very common (45). In a gastropod example, the rough periwinkle *Littorina saxatilis* has population-level genetic divergence at exceptionally small spatial scales (46). Because RNA-seq focuses on expressed coding regions and provides dense, high-quality single nucleotide polymorphism (SNP) data, using transcriptomics can more effectively distinguish genetic variation among individuals and populations. By integrating CRISPR-Cas9 gene manipulation with transcriptomes, variation in the coding sequences within the population being studied can be accommodated or targeted directly, facilitating precise functional examination of SNPs or large inter-individual genome variations. Consequently, transcriptomes can provide a practical foundation for gene-editing approaches in cases where genome assemblies remain incomplete or lack robust annotation, or where significant inter-individual genomic variation is known to occur.

In this paper we aimed to: (i) Develop an accessible pipeline for designing exon-specific gDNA primers and sgRNAs for CRISPR-Cas9 KO using a *de novo* transcriptome, (ii) generate an F0 *pax6* mutant in *C. fornicata* and confirm genome modification via both genotype and phenotype analysis and, (iii) propose *C. fornicata* as a comparative embryological system to study mollusc eye development and evolution.

## Methods

### Histological description of larval and adult eyes

Whole adult and larval *C. fornicata* (n = at least 5 per stage) were relaxed using drops of a saturated chlorobutanol solution (47). Once fully relaxed and not responsive to touch, animals were fixed in 4% paraformaldehyde (PFA) diluted in 0.2 µm filtered seawater at 4 °C overnight with gentle rocking. PFA was removed by rinsing three times (15 mins each on rocker at room temperature) in Phosphate Buffered Saline (PBS) before dehydrating to 100 % ethanol for storage at -20 °C. Tissues were cleared using histosol and embedded in paraffin wax (48). Wax blocks were serially sectioned at 5 µm on a Leica RM2125 rotary microtome, mounted on glass slides and rehydrated through a graded ethanol series. Slides were stained with haematoxylin and eosin (H&E) and imaged on a light microscope (Zeiss Axioscope A1 with Zen Blue software and a Zeiss Axiocam 503 colour camera).

### Transcriptome assembly

#### Short reads

Adult *C. fornicata* were obtained from the Marine Resource Centre at the Marine Biological Laboratory (Woods Hole, MA, USA) in July 2019 and maintained under standard conditions (49). Adults were shucked to extract embryos, which were staged according to previous work (24). Collected stages included ovoid (130 hpf), early organogenesis (170 hpf), late organogenesis (196 hpf), early veliger (228 hpf), late veliger (288 hpf), crawling juveniles, and adult mantle tissue. One capsule of approximately 200 full-sibling embryos were pooled to make a single sample and four samples were collected per stage. Samples were snap frozen in liquid nitrogen and stored at –80 °C. Total RNA was extracted from each sample using the RNAqueous-Micro Total RNA Isolation Kit, and libraries were prepared using Smart-seq2 with 10 cycles of cDNA amplification. Barcoded libraries were pooled and sequenced on an Illumina NovaSeq 6000 (paired-end, 150 bp). Raw reads were quality-checked using using FastQC (v0.11.9) and trimmed, using TrimGalore (v0.6.6) (parameter = --nextera). TrimGalore was also used to trim reads for quality and length (parameters = --length 70, --q 30). rRNA contamination was checked using Bowtie2 (version 2.4.2) and removed where present in libraries.

#### Long reads

Late organogenesis (196 hpf, two samples prepared as above), head vesicle (210 hpf, five samples), early veliger (228 hpf, one sample), late veliger (288 hpf, one sample), and adult mantle tissue (one sample) were used for long-read sequencing. RNA was extracted using phenol-chloroform and samples were diluted to 100 ng/µL prior to being combined in equal parts. 1 µg of RNA was sent to Novogene for isoform sequencing (PacBio sequel II). Long reads were processed using the Iso-seq pipeline, rRNA-filtered, and clustered to produce full-length transcripts. *C. fornicata* full-length RNA Circular Consensus Sequences (CCS) were generated by Novogene (79,255 reads). Read demultiplexing and the removal of polyA tails and library-prep-primers (68,396 reads) were conducted using lima (version 2.7.1; lima --isoseq --peek-guess) and IsoSeq (version 4.0.0; isoseq3 refine --require-polya). Full-length transcripts were *de novo* clustered and polished using Isoseq (version 4.0.0; isoseq3 cluster --use-qvs) (number of reads remaining: 5,645).

#### Assembly

The *C. fornicata* transcriptome was assembled using SPAdes (version 3.15.4; rnaspades.py -- pacbio) combining both the trimmed short-read and trimmed long-reads to produce a high coverage transcriptome (50). Redundancy was reduced using TransDecoder to retain coding sequences (51) and CD-HIT to cluster highly similar transcripts (37). The resulting assembly comprised 95,319 transcripts with an N50 of 1,702 bp and 83% BUSCO completeness, representing a substantial improvement over a short-read-only assembly and forming the consensus reference transcriptome used for all subsequent analyses (see *Data Availability* for transcriptome access).

### Candidate gene identification and alignment for putative gene intron/exon structure

A single *pax6* sequence was identified in the *C. fornicata de novo* transcriptome using a reciprocal BLASTN strategy (53). Publicly available *pax6* mRNA sequences from caenogastropods were downloaded, the *C. fornicata* transcriptome was used as a database and the caenogastropod *pax6* mRNA sequences were used as queries. The resulting single putative *pax6* gene in the *C. fornicata* transcriptome was then used as a query to search the Nucleotide collection (nt) database to confirm the top hits were all *pax6* genes. To further confirm that the extracted *C. fornicata* sequence was a true *pax6* homolog, phylogenetic relationships were analysed in Geneious Prime. Seven molluscan *pax6* mRNA sequences (*Gigantopelta aegis,* XM_041510489.1; *Crassostrea virginica,* XM_022437898.1; *Pecten maximus,* XM_033896327.1; *Physella acuta,* XM_059292525.1; *Biomphalaria glabrata,* XM_056044551.1; *Pomacea canaliculata,* XM_025231455.1; and *Littorina saxatilis,* XM_070344187.1), one polychaete worm (*Owenia fusiformis,* KY809737.1_cds_AUN27663.1_1), a cnidarian *Acropora millepora* (XM_029356362.2), and a fruit fly *Drosophila melanogaster* (NM_001272151.1), *twin of eyeless* mRNA were aligned using Clustal-Omega (54). Sequences from *A. millepora* and *D. melanogaster* were included as outgroups. A neighbour-joining consensus tree was inferred using the Tamura-Nei (TN93) nucleotide substitution model, which accounts for unequal base frequencies and different rates of transitions versus transversions, providing a realistic model for cross-species sequence comparisons (55). Branch support was assessed with 500 bootstrap replicates, a number sufficient to generate statistically robust values, and a 70% bootstrap threshold was applied to highlight strongly supported nodes (56,57). Trees were visualized with branch lengths and bootstrap values displayed (*Figure S1*).

To identify the putative genomic intron/exon structure of the *pax6* gene in *C. fornicata*, the *cf-pax6* mRNA sequence was aligned, using NCBI BLAST discontinuous megablast for more dissimilar sequences (58), to both genomic DNA (gDNA) and mRNA of other gastropod species *pax6* genes. Species used included the rough periwinkle, *Littorina saxatilis* (XM_073044187.1), the bladder snail, *Physella acuta* (XM_059202525.1), the scaly-foot snail, *Gigantopelta aegis* (XM_041510489.1), and the apple snail, *Pomacea canaliculata* (XM_025231455.1). Protein domain structure for *cf-pax6* was inferred using *L. saxatilis pax6* mRNA and InterProScan (59).

### Primer design and testing for gDNA PCR amplification

Using the putatively conserved intron/exon structure from the genomic alignments, primer design was focused solely within a putative exon sequence, ensuring primer pairs did not span multiple exons or exon/intron boundaries. Exon three was chosen as a target exon due to its proximity to the start of the homeodomain, which has a role for eye development in other models (60,61). Primers were designed using NCBI Primer-BLAST, with *C. fornicata de novo* assembled reference transcriptome as the custom reference database (62). Primer-Blast uses the Primer3 software coupled with BLAST to identify potential primers for GC content, annealing temperature, and dimer formation while searching for off-targets against the reference transcriptome (62). Primer pairs were chosen that had the least off-target sequences (<2 with >2 mismatches within off-targets). A limitation of not having a genome is that primers could not be tested *in silico* for off-target binding in the genome. Primer sequences were ordered from Integrated DNA Technologies (IDT) as desalted stocks and were rehydrated to 100 µM concentration using nuclease-free water.

Primers were tested using gDNA extracted from fresh adult foot tissue with the E.Z.N.A. Mollusc and Insect DNA Kit (Omega Bio-tek, D3373). A touchdown PCR protocol was employed to improve primer specificity. The DreamTaq 2X Master Mix kit was used (ThermoFisher, Product No. 11816843) in 25 µL reactions (95 °C 30 sec, [95 °C 30 sec, T_M_ + 10 °C 30 sec, 72 °C 1 min]x10, [9 5°C 30 sec, last T_M_-1 30 sec, 72 °C 1 min] x2 until at original T_M_, 72 °C 10 min) with an annealing temperature determined by the New England Biolabs Tm Calculator for Taq DNA Polymerase. PCR products were run on a 1.5 % agarose gel. Primer pairs that produced a single band at the expected amplicon of size were cleaned using the QIAquick PCR Purification Kit (Qiagen, #28104) and sent for Sanger sequencing to confirm successful amplification of the target exon sequence. By only selecting primer pairs that produce a single PCR band the issue of primer specificity and off-target amplification was overcome.

### Designing single-guide RNAs (sgRNAs)

Methods for sgRNA design were based on previous literature with modifications to account for using a transcriptome (16). CRISPRscan was used for initial sgRNA design; it assigns a gRNA score from 0-100 based on various factors such as GC content, nucleotide structure and presence of a protospacer adjacent motif (PAM) sequence (63). The target exon was input as a fasta file into CRISPRscan with “choose organism” set to “no search”, since there is no reference genome to test off-target sequences in *C. fornicata*. sgRNAs were filtered for a score >50. Filtered sgRNAs were next input into the IDT sgRNA design for further quality screening (16). IDT sgRNA design works similarly to CRISPRscan, and was used to provide a secondary quality screening for optimal sgRNA design (64). A score of 0-100 is assigned to each sgRNA, with 70 being considered a high-efficiency score. The species selected was “other” for no off-target screening. sgRNAs were further filtered with a cut-off score >40.

CRISPRscan and IDT screened candidate sgRNAs were next screened *in silico* for off-targets using blast-plus v2.12.0 (65) and the *C. fornicata* reference transcriptome. Candidate sgRNAs were concatenated into a multi fasta file and BLASTN, using --task blast-short function, was used to find potential off-target sequences in the transcriptome. sgRNAs that had less than 2 mismatches in their list of off-targets were removed. As with primer design, a limitation of not having a genome is that sgRNAs could not be tested for off-target binding to non-coding regions of the genome.

sgRNAs that had a CRISPRscan score of >50, an IDT score of >40, and >2 mismatches in any off-targets hits were kept for further screening (*Table 2*). sgRNAs were ordered from Synthego. sgRNAs were resuspended in nuclease-free water to a concentration of 100 µM and stored at -20 °C for short term usage. Cas9 protein was also ordered from Synthego and stored at -20 °C for short term usage (SpCas9 2NLS nuclease, 300 pmol, Synthego, USA). sgRNAs and Cas9 protein were stored at -80 °C for long term storage.

**Table 1.**
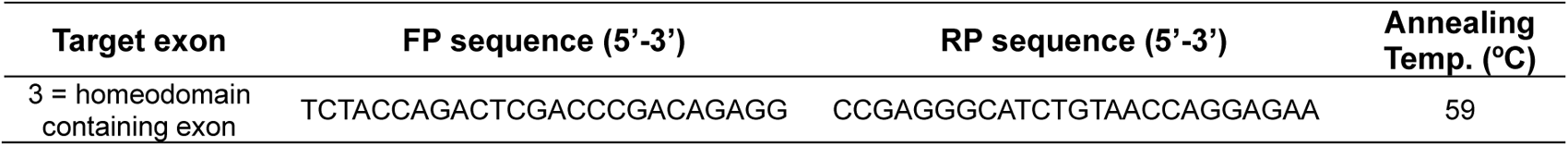
gDNA primers for *cf-pax6* target exon.

**Table 2.**
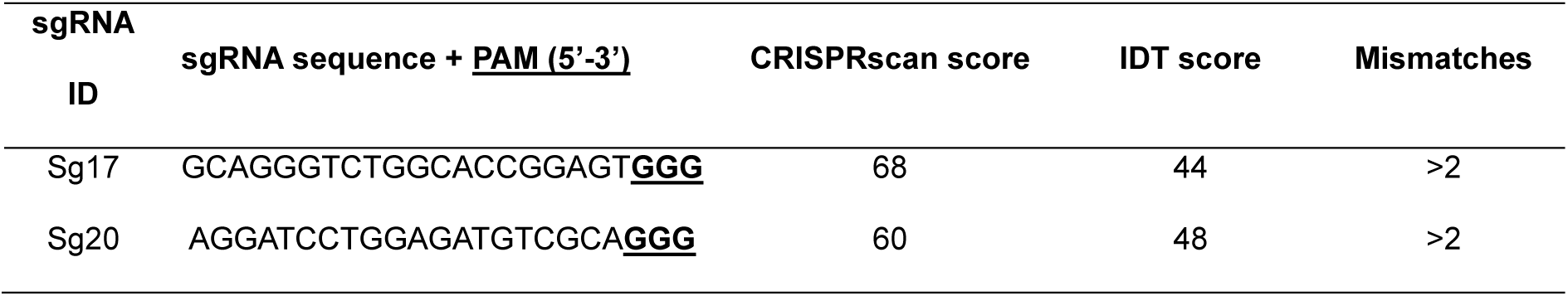
Single-guide RNA sequences used for *cf-pax6* Kos.

### Embryo harvest and cultivation for KO experiments

Adult broodstock snails were obtained from Marine Resources Centre (MRC) at the Marine Biological Laboratory in Woods Hole, Massachusetts, USA. The MRC collected subtidal adult stacks via trawling in late winter/early spring and maintained them in chilled flow through aquaria (≤15 °C) to dissuade spawning until required for experiments. To obtain one-celled embryos, adult stacks were placed in a warm >18 °C flow through aquaria overnight (22). Adults were gently shucked in the morning using an oyster shucking knife. Embryo capsules were removed by pinching the single joining point of their surface attachment before placing them in a dish of 0.2 µm-filtered sea water (FSW). Adults were gently pieced back together and left for several hours upside down to reattach to their stacks; embryos were transferred to the laboratory for microinjection.

Embryos were staged using a dissection microscope and capsules containing one-celled zygotes were vigorously rinsed using a household spray bottle filled with FSW to reduce ciliates and other unwanted contaminating microorganisms (19,24). Vigorous rinsing was repeated 3-4x with excess water removed in between. Cleaned capsules were opened and embryos were removed and sorted for microinjection, or directly reared on gelatin-coated dishes as uninjected controls (0.01 % Knox gelatine, 0.04 % of 32 % PFA, in 50 mL FSW and dried at room temp. overnight), in 1X Penicillin-Streptomycin FSW (5X stock PenStrep FSW = 0.05 g Streptomycin sulphate salt, Sigma Aldrich S9137-25G, 0.0125 g Penicillin G sodium salt, Sigma Aldrich P3032-1MU, in 50 mL, diluted 1:5 with FSW to make working 1x solution (20,22)). The gelatin-coating prevented embryos from sticking to the plastic dish. Embryo dishes were kept in a “hydration chamber” to prevent the evaporation of seawater increasing salinity (a large 120 mm petri dish, lined with deionized water-soaked paper towels with the lid on). All embryos were reared in an incubator at 18 °C. 1X PenStrep FSW was changed, embryos were scored, and cleaned of debris and mortalities daily. Embryos were carefully transferred to new dishes as needed if there was a build-up of debris. Embryos were reared until desired stage depending on experiment.

### Microinjection

Single-cell embryos were used for CRISPR KO experiments. All controls are full siblings to the experimental injected embryos and were from the same capsule. Staged and cleaned embryos were carefully transferred to a microinjection dish using a mouth pipette and gelatine coated, hand-pulled glass Pasteur pipette. Microinjection dishes were filled halfway with 2% agarose in FSW and imprinted with a custom 3D-printed stamp with 150 µm wells *(see Data Availability for 3D file, Figure 1a)*; once set, dishes were filled with 1x PenStrep FSW. Embryos were moved into imprinted microinjection lines in the dish using an eyelash mounted to a Pasteur pipette (*Figure 1b*).

**Figure 1.**
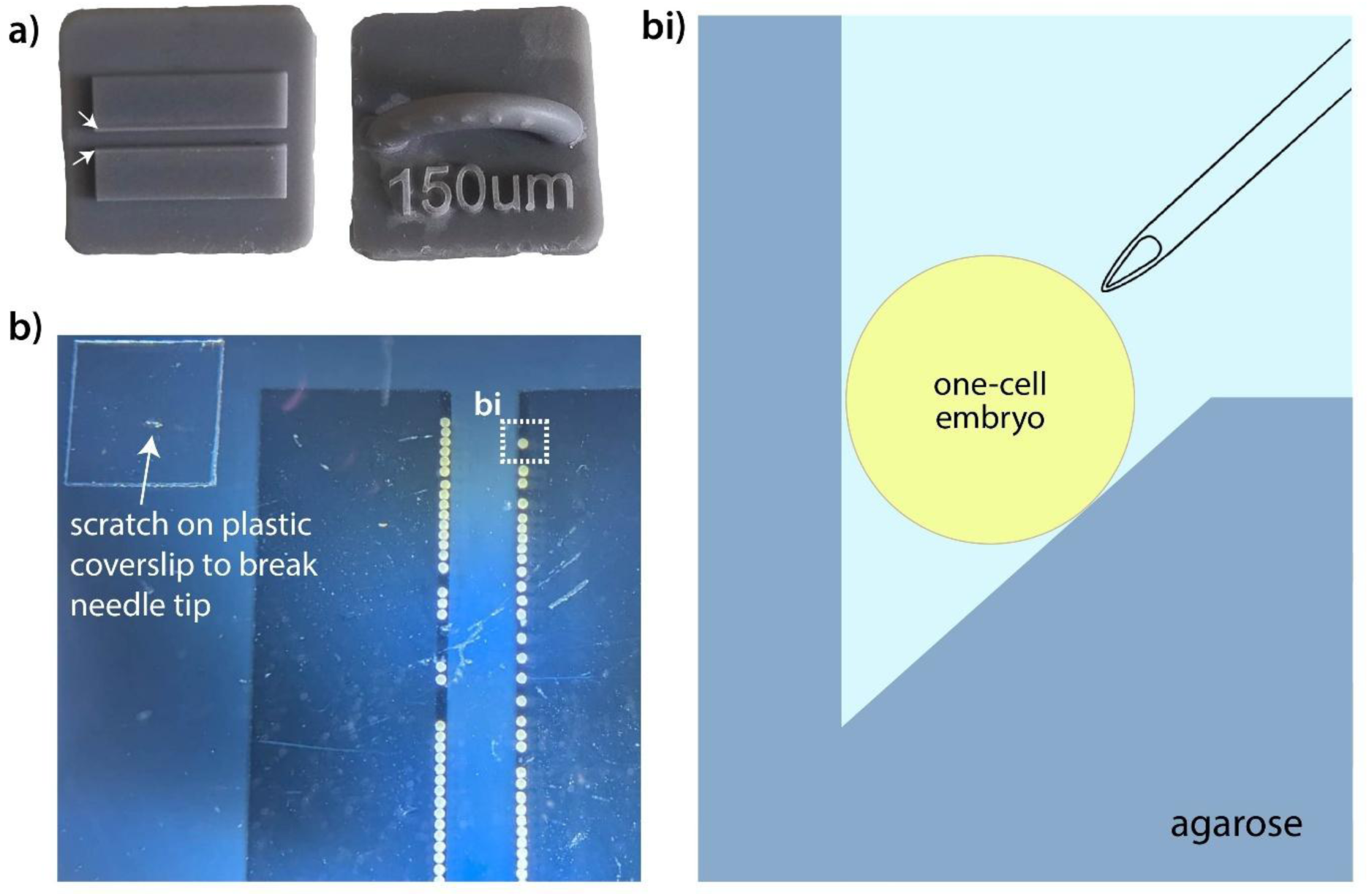
3D printed mould for embryo microinjection. *(a)* 3D printed mould to create 150 µm wide channels in agarose (channels highlighted with white arrow heads)*. (b)* Imprinted agarose dish with one-celled *C. fornicata* embryos lined up in 150 µm channels, viewed under a dissecting microscope, plastic coverslip used to break the needle also in view. *(bi)* Schematized cross section of the imprinted microinjection dish with embryo in microinjection channel.

Glass capillaries were pulled for microinjection using a Sutter P-1000 micropipette puller with a 2.5 mm x 2.5 mm box-type filament and thin-wall capillary tubes with a 1 mm diameter (World Precision Instruments, TW100-4). Microinjection needles were backloaded with the microinjection mixture. Fluoro-Ruby was used to confirm success of microinjection and was usually visible until the veliger stage (Invitrogen™ Dextran, Tetramethylrhodamine, 10,000 MW, Lysine Fixable, D1817). A Leica M165 FC dissecting scope was used to visualize embryos, with a Picospritzer II (General Valve Corporation) and an MMO-4 Three-Axis Hanging Joystick Oil Hydraulic micromanipulator from Narishige to microinject embryos. Pressure settings were around 1300 psi. The needle was gently broken by tapping it against a scratch in a plastic coverslip *(Figure 1b)*, until a bead of liquid (visualized in pink because of the dextran dye) about 5% the size of an embryo was pulsed out.

### Ribonucleoprotein (RNP) complex mixture

sgRNA and Cas9 concentrations in spiralian microinjection CRISPR-Cas9 experiments were reviewed (*Table 3*). The concentration selected for our KO experiments were based on recent success in *C. atrasolea* as it was the most closely related species and was in the middle of the range of concentrations reported (*Table 3*). For single-guide injections, the RNP mix consisted of 1.33 µL of 900 ng/µL stock sgRNA, 0.8 µL of 20 µM stock Cas9, 2 µL Fluoro-Ruby, and 3.87 µL nuclease-free water.

**Table 3.** sgRNA and Cas9 protein concentrations from CRISPR-Cas9 experiments in Spiralia using microinjection into one-celled zygote for delivery.

| Species | Approximate<br>zygote<br>diameter<br>(μm) | sgRNA<br>(ng/μL) | Cas9<br>(ng/μL) | Citation |
| --- | --- | --- | --- | --- |
| Mollusc, gastropod:<br>*Pomacea canaliculata* | 100 | 10 | 10 | (17) |
| Mollusc, gastropod:<br>*Lottia goshimai* | 130 | 280 | 1800 | (15) |
| Mollusc, gastropod:<br>*Lymnaea stagnalis* | 250-350 | 100 | 1500 | (18) |
| Mollusc, gastropod:<br>*Crepidula atrasolea* | 160 | 300 | 600 | (16) |
| Mollusc, gastropod:<br>*Crepidula fornicata* | 200 | 250 | 1800 | (12) |
| Mollusc, gastropod:<br>*Crepidula fornicata* | 200 | 300 | 600 | This paper |
| Mollusc, bivalve:<br>*Magallana gigas* | 50 | 500 | 500 | (14) |
| Mollusc, cephalopod:<br>*Doryteuthis pealeii* | 1000 | 1000 | 1000 | (13) |
| Annelid:<br>*Capitella teleta* | 100 | 125 | 2000 | (7) |
| Annelid:<br>*Platynereis dumerilii* | 160-180 | 20 | 300 | (66) |

### CRISPR KO experimental design

Microinjected embryos and sibling controls were raised to early ovoid for genotyping and cutting efficiency analysis and late organogenesis to early veliger for phenotyping (with some also phenotyped embryos also genotyped). For initial genotyping, and to test cutting efficiency of each sgRNA, ten embryos per sgRNA were fixed in 100 % ethanol and stored at -80 °C until genomic DNA (gDNA) extraction. Two wild type uninjected embryos and two no Cas9 injected controls were fixed per sgRNA.

For morphological analysis, embryos were fixed at late organogenesis to early veliger stages (200 - 228 hpf) in 4% PFA for one hour at room temperature and then rinsed 3x 15 minutes in 1x phosphate buffer saline (PBS) before moving through a dehydration gradient into 100 % ethanol. Samples were stored at -20 °C for long term storage. Embryos that died before the late organogenesis or early veliger stage were not collected or scored for phenotyping because the eyes had not yet developed.

### Single embryo gDNA extraction and PCR amplification for genotyping

To genotype crispant KO embryos and sibling controls, sufficient quality and quantity of gDNA was required from a single embryo. DNA was extracted with the Monarch Spin gDNA extraction kit, with minor modifications (New England Biolabs, #T3010). Briefly, storage ethanol was pipetted off the embryo and a Kimwipe was used to remove any residual ethanol. 100 µL Lysis Buffer with 20 µL 20 mg/mL Proteinase K from the kit was immediately added, such that the embryo did not completely dry out, and incubated at 65 °C for 5 minutes. 400 µL Binding Buffer with 4 µL carrier RNA was added to the samples, which were then vortexed until homogenized (Ambion® Carrier RNA 1 mg/mL, ThermoFisher #4382878). Bind and wash steps were carried out per the kit’s instructions. For elution, columns were added to a new 2 mL collection tube, and 25 µL of pre-warmed at 60 °C elution buffer was added to each. This was left to sit for 2 minutes at room temperature before spinning at 13000 rpm for 1 minute. Extracted samples were assessed for quantity and purity on a Nanodrop Lite Plus (ThermoFisher) and stored at -20 °C for long-term storage.

For genotyping, samples were amplified using a touchdown PCR protocol in 50 µL reactions using the 2X DreamTaq Master Mix (25 µL Master Mix, 6 µL Nuclease Free H_2_O, 2.5 µL 5 µM Forward Primer, 2.5 µL 5 µM Reverse Primer, 2 µL 4 mM Magnesium Chloride, 4 µL gDNA). A touchdown PCR protocol was chosen because it can increase primer specificity (67). Positive and negative controls were included for each PCR, with positive control gDNA contribution from adult mantle tissue extracted using the same methodology as above. Post-PCR, samples were cleaned using the QIAquick PCR Purification kit and eluted into 30 µL elution buffer. Cleaned PCR samples were stored at -20 °C for long-term storage or used directly into another touchdown PCR protocol following the same specifications as above to increase concentration of product. Following the second PCR, 10 µL of each sample ran on a 1.5% agarose gel with SYBR Safe (Invitrogen SYBR Safe DNA Gel Stain, S33102) in 1x Tris-acetate-EDTA (TAE) buffer at 70V for 45 minutes with a 100 bp ladder for size assessment. 5 µL of each sample was also run on a high-sensitivity Agilent TapeStation. Samples with bands of an appropriate amplicon size were cleaned once more using the QIAquick PCR Purification kit and nanodropped for concentration and quality assessment.

### Sanger Sequencing and ICE analysis

Each PCR product was sent to Eurofins Genomics (Cologne, Germany, 20 ng per PCR product) for Sanger sequencing. To determine CRISPR-Cas9 cutting, chromatograms were examined for signal disruption at the site of the sgRNA. Additionally, samples were submitted to an Interference of CRISPR Edits (ICE) analysis by Synthego, which parses out discordant peaks and quantifies the different mutations within a single embryo by comparing a wild type chromatogram to a mutant chromatogram (68). ICE analysis provides a proxy for sgRNA efficiency and whether mutations are out-of-frame or in-frame.

### Phenotyping CRISPR-Cas9 mutants

Samples stored in 100 % ethanol were examined under a Leica M165 FC dissecting microscope for morphological comparison. Embryos were categorized by having no eyes, one eye, or two eyes, and sample numbers were totalled for each sgRNA treatment. Some examples of live embryos were also imaged prior to fixation on Leica M165 FC dissecting microscope fitted with a Leica DFC7000 camera.

## Results

### *Crepidula fornicata* have simple cup larval eyes that develop into lenticular adult eyes

During the organogenesis stage two black pigmented anterior eye spots become clearly visible in *C. fornicata* embryos and so we sought to histologically characterise them from organogenesis onwards. Sections of the developing eyes were obtained in both sagittal and transverse planes. At late organogenesis, the larval eyes are simple cups that appear to form via invagination of the surface ectoderm *(Figure 2a-c)*. The larval eyes are covered by a thin epithelium, the early cornea, that closed the cup structure *(Figure 2ai, bi, bii)*. The cup structure consisted of a one-cell thick layer of sensory cells, some of which contained black pigment granules *(Figure 2ai, bi, bii)*. The one-cell thick cup structure remained into the veliger stage, the amount of pigment granules increased, and the small lens became refractive *(Figure 2c, ci)*.

**Figure 2.**
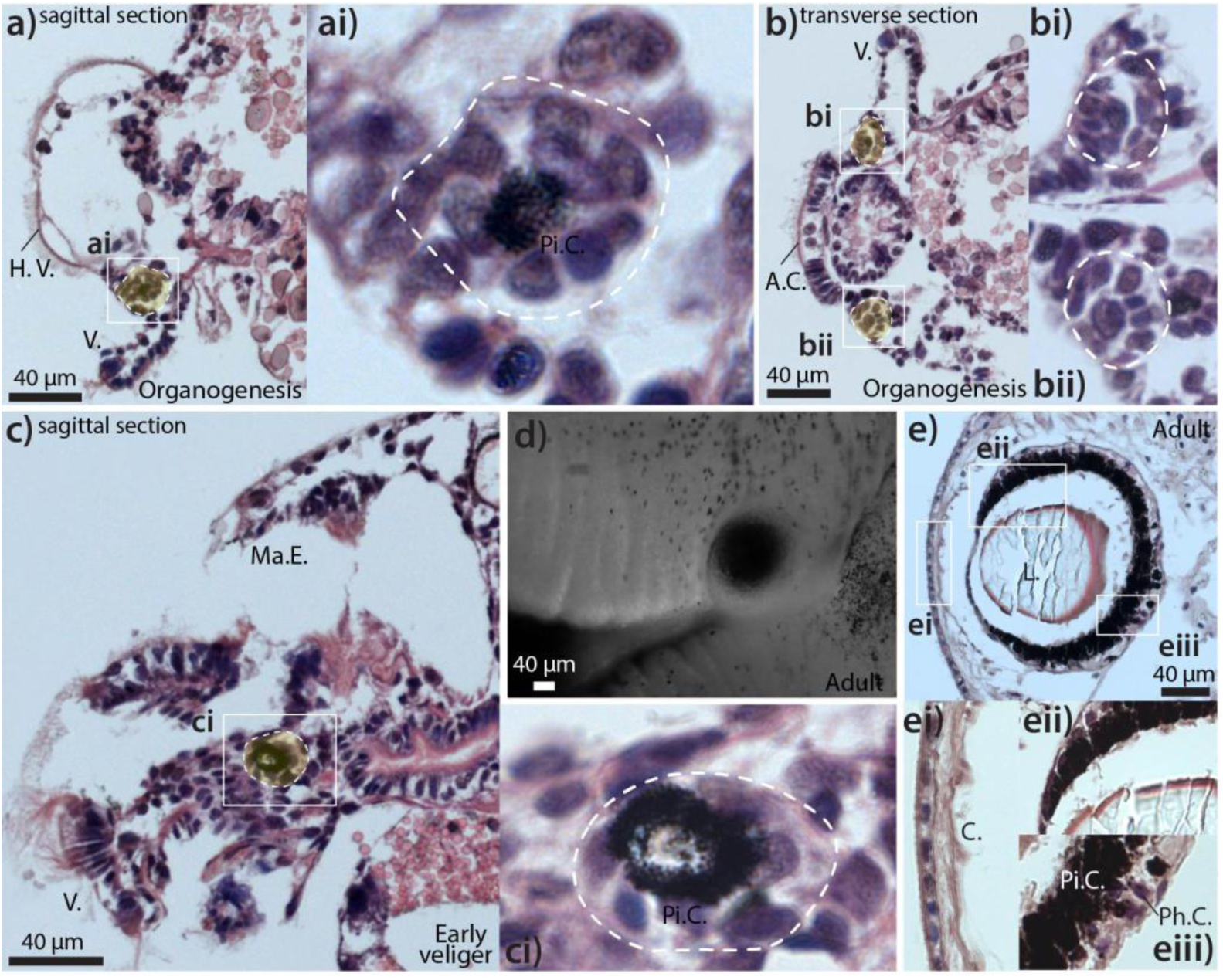
H&E histology of *C. fornicata* larval and adult eyes. *(a)* Sagittal section of late organogenesis stage embryo, simple cup eye highlighted in yellow. *(ai)* Zoom-in from white box highlighted in (a) showing arrangement of pigment cell and putative sensory cells covered by an epithelium. *(b)* Transverse section of late organogenesis stage embryo, two simple cup eyes visible in section plane, highlighted in yellow. *(bi & bii)* Zoom-in from white boxes highlighted in (b) showing cellular arrangement of both eye cups covered by an epithelium. *(c)* Sagittal section of early veliger stage embryo, simple cup eye with increased pigmentation highlighted in yellow. *(ci)* Zoom-in from white box highlighted in (c) showing arrangement of pigment cells and putative sensory cells covered by an epithelium. *(d)* Wholemount image of fixed adult eye at base of cephalic tentacle. *(e)* Cross section of lenticular adult eye. *(ei)* Zoom-in from white box highlighted in (e) showing cornea. *(eii)* Zoom-in from white box highlighted in (e) showing transition to non-sensory area of retina and space between the retina and lens. *(eiii)* Zoom-in from white box highlighted in (e) showing sensory cells in the retina including a layer of pigment cells and photoreceptor cells. H. V. = Head Vesicle; V. = Velum; Pi. C. = Pigment Cell; A. C. = Apical Cell Plate; Ma. E. = Mantle Edge; L. = Lens; C. = Cornea; Ph. C. = Photoreceptor Cell.

As adults, *C. fornicata* have clearly visible black spherical eyes at the base of both of their cephalic tentacles *(Figure 2d)*. In large, likely fully grown, female specimens the diameter of the eye was approximately 160 µm *(Figure 2d,e)*. Histological analysis revealed that adult *C. fornicata* have lenticular eyes including a layer of cuboidal epithelial cells in the cornea, an acellular lens that did not appear to be in direct contact with the cornea, and a retina *(Figure 2e, ei, eii)*. The retina had a dense layer of pigment cells, followed by a thinner layer of sensory photoreceptor cells *(Figure 2eiii)*.

### Successful design of exon-specific primers and sgRNAs for CRISPR-Cas9 without a genome

In the phylogenetic analysis, the *cf-pax6* transcript was clustered most closely to other gastropods sequences such as *P. canaliculata* and *L. saxatilis*; all nodes in the resulting tree were supported by bootstrap values greater than 87, indicating strong confidence in the branching relationships (*Figure S1*). *C. fornicata pax6* mRNA was aligned to the gDNA of four species of gastropods to determine the putative intron-exon structure of the *cf-pax6* gene (*Figure 3*). The five gastropod species had a largely conserved *pax6* gene structure, with 10-12 exons and 70-80 kbp in length (*Figure 3*). The *cf-pax6* gene from our *de novo* assembled reference transcriptome is a partial transcript that spans the middle portion of *pax6* gene (*Figure 3*). *cf-pax6* aligns with exons 2, 3 and 4 of the *L. saxatilis pax6* gDNA (*Figure 3a*). When aligned with *P. acuta pax6* gDNA, *cf-pax6* aligned to exons 4, 5, and 6 (*Figure 3b*). In both *G. aegis* and *P. canaliculata* alignments, *cf-pax6* aligned to the middle of the gene, exons 4 and 5, and 5 and 6, respectively (*Figure 3c, d*).

**Figure 3.**
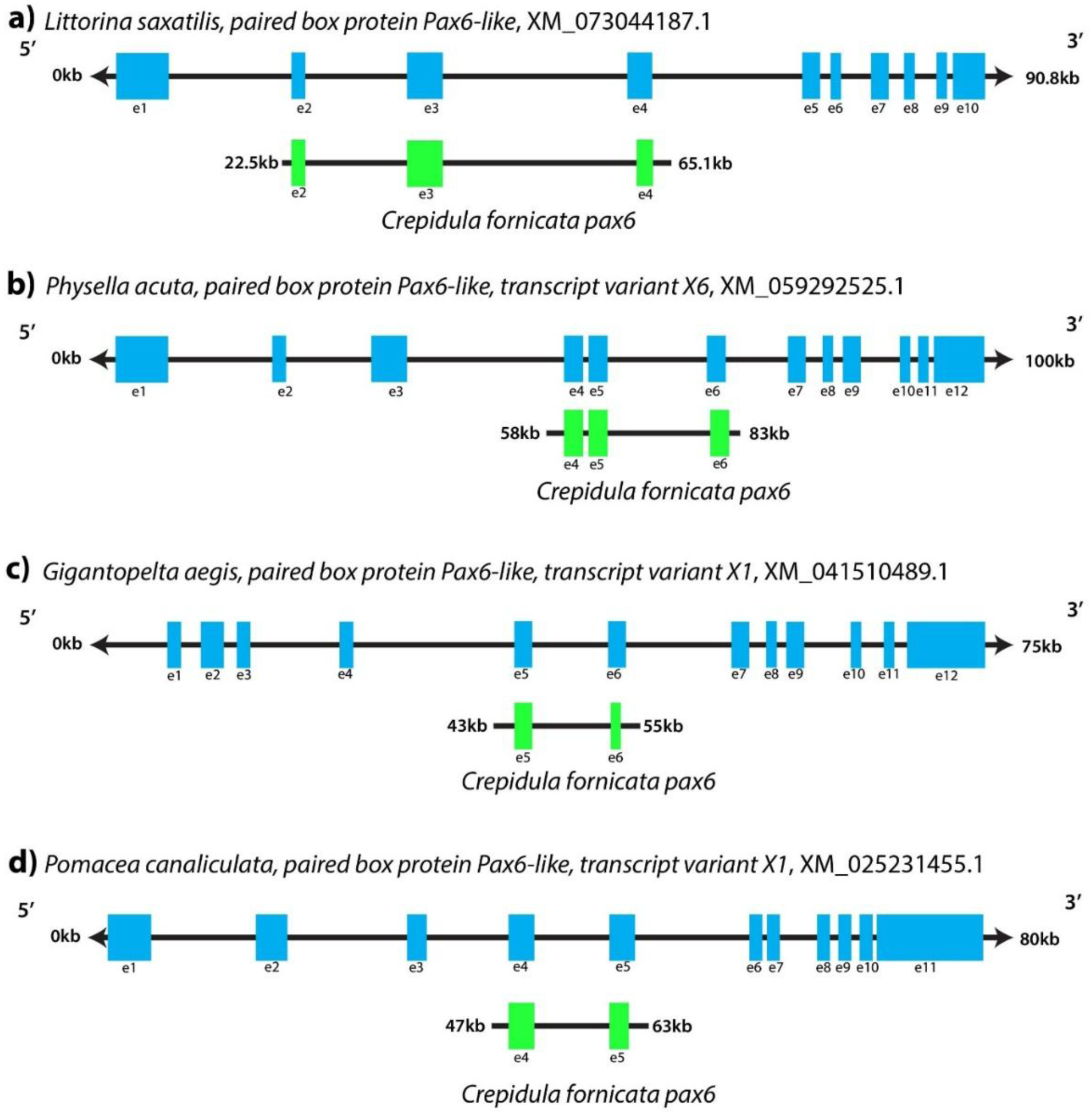
*cf-pax6* mRNA partial sequence aligned to the *pax6* gDNA of other gastropods. (*a-d*) Blue boxes represent exons for the reference gastropod’s *pax6* gDNA. Green boxes represent aligned regions, and therefore proposed exons for *cf-pax6*. (*a*) Alignment of *Littorina saxatilis pax6* gDNA to *cf-pax6* mRNA. (*b*) Alignment of *Physella acuta pax6* gDNA to *cf-pax6* mRNA. (*c*) Alignment of *Gigantopelta aegis pax6* gDNA to *cf-pax6* mRNA. (*d*) Alignment of *Pomacea canaliculata pax6* gDNA to *cf-pax6* mRNA.

*L. saxatilis* was the most closely aligned species to *cf-pax6* (based on BLASTN e-value, 2e^-135^) and was used as the putative reference for intron/exon structure in *cf-pax6* (*Figure 4a*). Two sgRNAs were designed within the *cf-pax6* partial sequence in exon three, within the homeodomain (*Figure 4a*). Primers were successfully designed for *cf-pax6* exon 3 (*Table 3, Figure 4b*).

**Figure 4.**
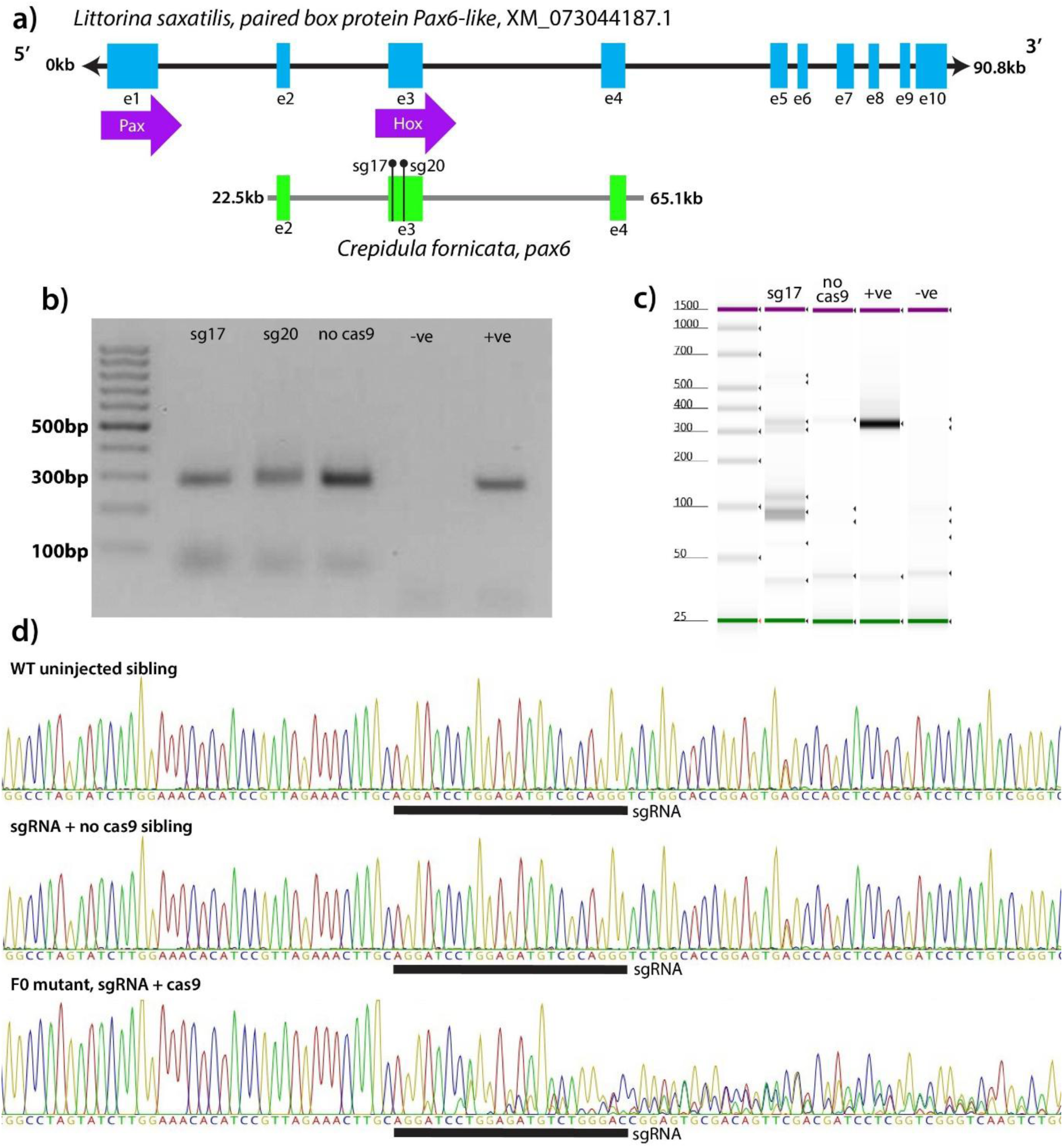
Genotyping *cf-pax6* mutant F0 embryos. (*a*) Alignment of *L. saxatilis pax6* gDNA with *cf-pax6* partial mRNA sequence. Black vertical lines capped in a ball represent sgRNA placement in *cf-pax6*. Purple arrows represent protein domains for *L. saxatilis pax6*. (*b*) An agarose gel showing PCR product amplified from single embryos. (*c*) A TapeStation digital gel showing experimental samples and controls for *cf-pax6* KO experiments. (*d*) Chromatograms displaying wild type uninjected sequence (top), wild type sgRNA only control with no Cas9 (middle) and mutant sgRNA + Cas9 *cf-pax6* F0 (bottom). All embryos were siblings from the same capsule.

### Successful genotyping *cf-pax6* F0 CRISPR mutants

gDNA was extracted from single microinjected embryos and the target *pax6* exon was successfully amplified and clearly visible on an agarose gel with a desired amplicon size of 288 bp (*Figure 4b*). The same PCR products run on an Agilent TapeStation contained the identical 288 bp band (*Figure 4c*). Mutant KO samples run on a high-sensitivity Agilent TapeStation allowed the visualization of genetic mosaicism as multiple bands of different sizes were also visible (absent in the wild type samples, *Figure 4c*). Mosaicism was therefore visible on a high-sensitivity Agilent TapeStation tape but otherwise not detectable through standard agarose gel.

Of the 5 samples genotyped for both sgRNA17 and sgRNA20, sgRNA17 cut 60 % (3/6) of the time, and sgRNA20 cut 80 % of the time (4/5). Both the uninjected wild type controls and sgRNA with no Cas9 controls had clean chromatograms that did not indicate any mutations (*Figure 4d*). All samples where the sgRNA has successfully cut displayed discordant, multiple peaks at the site of the sgRNA, compared to the clean and singular peaks of the controls (*Figure 4d*). ICE analysis was used to determine the range of mutations in a single mosaic F0 crispant embryo. For sgRNA 17, a single embryo had 68 % of the *cf-pax6* copies wild type, 10 % had a +1 bp mutation, and 8 % had a -6 bp mutation (*Table S1*). For sgRNA 20, a single embryo contained 24 % of wild type *cf-pax6* copies, with 28 % -8 bp and 23 % -19 bp mutations (*Table S1*). For both sgRNAs, other in- and out-of-frame mutations made up the remaining percentages of the copies of *cf-pax6* (*Table S1*).

### Phenotyping *cf-pax6* F0 CRISPR mutants

As the eyes develop by the late organogenesis stage and are easily visible with no stains or dyes, embryos were photographed (either alive, or after fixation and storage in ethanol) to determine and quantify eye phenotypes. Embryos with both eyes had otherwise normal morphology (*Figure 5a*). For the one-eyed embryos, the side of the embryo without an eye appeared morphologically identical to the side with an eye (*Figure 5b*). Embryos with no eyes were grossly morphologically the same as their sgRNA only-injected sibling controls (*Figure 5c*), except some mutant embryos also exhibited smaller heads and velum. For both sgRNA17 and sgRNA20, a proportion of microinjected embryos had a phenotype of one or no eyes (*Figure 5d)*. Only eye phenotypes were quantified in the present study. There were developmental timing differences between the mutant and wild type or no-Cas9 control counterparts, embryos injected with Cas9 and sgRNA had an approximate 24-hour lag in development behind their no-Cas9 control or uninjected siblings.

**Figure 5.**
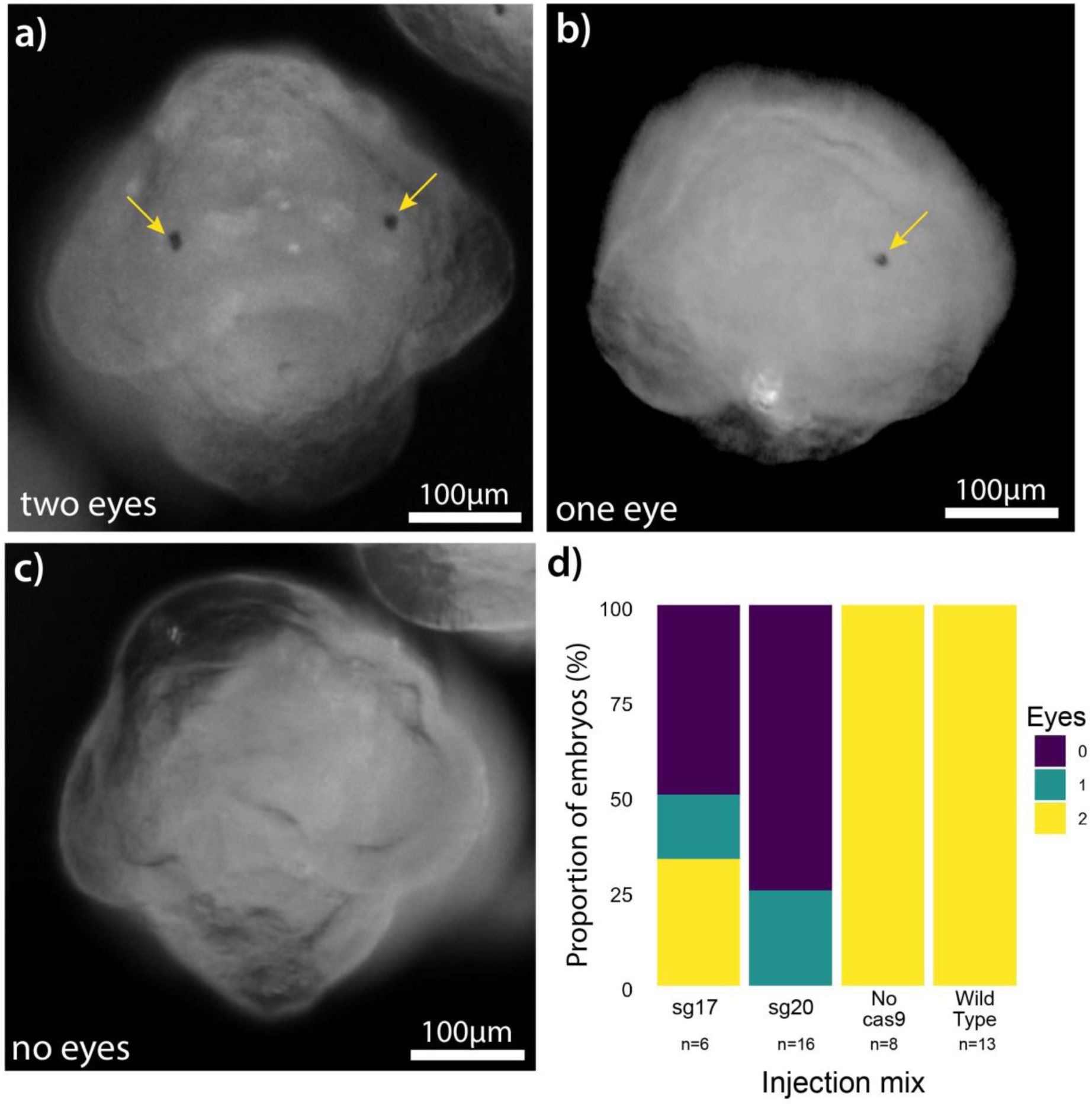
Phenotyping *cf-pax6* mutant F0 embryos. (*a-c*) Anterior views of organogenesis stage *C. fornicata* embryos injected with sgRNA20 for *cf-pax6*, scale bar = 100 µm. All embryos are siblings. Yellow arrows indicate eyes. (*a*) embryo with two eyes (live image). (*b*) embryo with one eye (fixed specimen). (*c*) embryo with no eyes (live specimen). (*d*) Percentage of surviving embryos at the organogenesis stage with 0, 1, or 2 eyes. Total number of surviving embryos in each treatment are displayed below each bar.

For embryos injected with sgRNA17, 30 % of embryos had both eyes, 20 % had one eye, and 50 % had no eyes (*Figure 5d*). For embryos injected with sgRNA20, 25 % of embryos had one eye and 75 % of embryos had no eyes (*Figure 5d*). All embryos microinjected with an sgRNA but no Cas9, or their uninjected wild type siblings, had normal morphologies and developed two eyes (*Figure 5d*). All measurements were of embryos that survived to late organogenesis or early veliger stage (matched on developmental stage landmarks not time to account for the developmental delay). Embryos that died pre-collection were not genotyped or phenotyped.

## Discussion

The goals of this paper were: (i) develop an accessible pipeline for designing exon-specific gDNA primers and sgRNAs for CRISPR-Cas9 KO using a *de novo* transcriptome, rather than a genome, (ii) generate F0 *pax6* KO mutants in *C. fornicata* confirmed via both genotype and phenotype and, (iii) propose *C. fornicata* as a comparative embryological system to study mollusc eye development and evolution. Using alignments with other closely related gastropod species and several quality assessment tools for sgRNA efficacy, F0 crispant KO *cf-pax6* mutants with an eyeless phenotype were generated. More specifically, sgRNAs placed immediately before the homeodomain in *cf-pax6* caused mosaic out-of-frame mutations true to an F0 mutant and inhibited the development of one or both eyes creating a classic eyeless phenotype. Generating crispants with observable F0 phenotypes using CRISPR-Cas9-mediated mutagenesis is significant as it demonstrates the feasibility of this approach to be applied to higher through-put gene function screening work. In addition, such work can be achieved without an assembled genome, demonstrating the feasibility of gene-editing approaches in non-traditional model organisms where only transcriptomic resources are available. It is, however, important to note that single copy or highly conserved genes are most likely to be successful with this approach, as related species with annotated genomes are necessary. We also histologically characterised both eye development and mature adult eyes in *C. fornicata,* laying the foundation for future comparative eye development and evolution studies.

### Spiralian and molluscan models for eye development and evolution

*pax6* has a conserved function across the Metazoa (69,70), including within Spiralia, as a transcriptional regulator in eye development (71–74). To date, eye reduction phenotypes in *pax6* (or related *toy/ec* orthologues) mutants have been recorded in Arthropoda (34,75), Mammalia (76), Amphibia (61), and Actinopterygii (35,77). However, genetically tractable models to study loss-of-function mutations in spiralian *pax6* have remained inaccessible until recently (17). In this study, we provide a second example of a CRISPR-Cas9 induced *pax6* mutant with loss-of-eye phenotypes in ceanogastropods and therefore confirm the conserved function of *pax6* in this subclass of molluscs. Multiple genetically enabled model systems how expands the scope for eye development and evolution studies.

Molluscs have a diverse range of eye types, including complex camera-type eyes (78,79), mirror eyes (80), and pit eyes (81). Within the caenogastropod subclass, multiple groups have been studied in the context of visual system evolution. For example, the evolution of very large eyes (82), loss of eyes (83), and the regeneration of complex camera-type eyes (17). Here, we histologically described the development of a closed lenticular eye in the caenogastropod *C. fornicata*. During development the larval eyes were simple closed cup structures that included pigment cells, like those seen in other gastropod larvae (84–86). The adult eye had a cornea, acellular lens and multi-layered retina, although its resolution and sensitivity has yet to be determined. We also did not find evidence of a structure that could regulate the amount of light entering the eye and so it is unlikely to be the most complex camera-type eyes seen in other gastropods. Crucially, we have begun to characterise the development of the visual system in an embryologically, and now, genetically, enabled model system that can be used a comparative model. The generation of *cf-pax6* F0 crispant mutants therefore provides a platform for research into the evolution of eye diversity both within Mollusca but also across the Metazoa when coupled with other CRISPR mutant models (34,35,61,76).

In the gastropod *P. canaliculata*, *pax6* F2 null mutants where the pax and homeodomains were disrupted did not develop a retina or an eye stalk (17). Morpholino-driven knock-downs of the marine annelid, *C. teleta*, disrupted only the homeodomain, leaving the pax domain intact (60). Like the *cf-pax6* F0 mutant presented here, disruption of the homeodomain but not the pax domain was enough to inhibit the development of embryonic eyes in *C. teleta* (60). Domain-specific perturbations in *C. teleta* implicated separable functions for the paired and homeodomains (60). In vertebrates, a *pax6* crispant in the African clawed frog, *Xenopus laevis*, showed that disruption of the homeodomain with an intact pax domain inhibited eye development (61), mirroring the phenotypic results presented here. Taken together with previous results, our new data in *C. fornicata* suggests that specifically the homeodomain in the *pax6* gene is critical for the development of eyes.

### Molluscan models for molecular and developmental evolution

Molluscs have high levels of phenotypic and genotypic diversity making them a fantastic phylum for evolutionary research (11). Studying molecular and developmental evolution of gene function with emerging tools like CRISPR-Cas9 gene editing, however, is limited in molluscs by the availability of genomes and embryos suitable for manipulation via microinjection. Advancements have been made in developing techniques for the delivery of CRISPR-Cas9 into less robust embryos via electroporation in primary cell lines (8,9) and vitellotagging (10). However, accurate and specific sgRNA design remains limited by genome availability and genome variability between individuals (3–5). Molluscs present a unique challenge here, as they exhibit exceptionally high genetic diversity across multiple scales (43–46). Genome assemblies of the European flat oyster, *Ostrea edulis*, have shown extensive variation in genome structure and variant profiles among individuals, reflecting substantial standing genetic diversity (44). Comparative genomic analyses across 32 different mollusc species further highlight pronounced interspecies genomic differences, including expansions of gene families and highly heterozygous regions (43). Given this high level of genetic variation, approaches such as RNA sequencing, which focus on expressed coding regions and produce high-resolution SNP datasets, enable more precise discrimination of genetic differences within and between populations. Indeed, transcriptome-wide SNPs were used to resolve the origins of two independent *Mytilus* introductions into southeastern Australia, revealing that one originated from a Mediterranean *M. galloprovincialis* lineage and the other from an Atlantic lineage, and that both subsequently admixed with the native *M. planulatus* (87). This was previously undetected using genome-wide or few-locus approaches, and detection of this can affect the management strategies for invasive species (87).

Here, we establish a methodology for utilizing CRISPR-Cas9 in species without a genome, instead capitalizing on a cheaper and simpler *de novo* transcriptomes that can easily be generated at the individual or population level. Furthermore, this paper established that gene function in developmental contexts can be studied in an F0 mosaic crispant. Studying gene function in an F0 crispant KO is advantageous where rapid gene function screening is required, perhaps for projects studying multiple genes and when maintaining multiple generations is either not possible because of limitations in culturing (13), or impractical due to length of time to sexual maturity – as is the case in *C. fornicata* (20,22).

The advancement of CRISPR-Cas9 techniques in *C. fornicata* first established by Perry et al., (2015) and now developed further in this paper, opens new avenues to answer evolutionary questions in calyptraeid molluscs. A close relative of *C. fornicata*, *C. atrasolea*, is also a genetically enabled model for molluscan developmental biology but has a much smaller generation time (and brood size). *C. atrasolea* has established molecular biology techniques, an assembled genome (not yet publicly available) and CRISPR-Cas9 mediated KO (16,25,88). With development of an F0 *cf-pax6* mutant here and the F0 *C. atrasolea SMP1* mutant developed in *C. atrasolea*, CRISPR-Cas9 can now be used to answer evolutionary questions between closely related species within a gastropod genus (16), opening the door for powerful micro-evolutionary comparisons that could elucidate molluscan morphological evolution. Moreover, *C. fornicata* possesses different developmental and physiological traits to *C. atrasolea* and thus providing an excellent comparative system. For example, *C. fornicata* has increased ocean acidification resistance (89), indirect larval development (19), and is a prolific invasive species (90).

## Limitations

Lack of an annotated genome is not the only factor limiting the use of gene-editing in Spiralia. While this paper has established the feasibility of gene-editing with only a *de novo* transcriptome, animal husbandry is a major inhibitor. To achieve gene-editing in single cell embryos, embryos must be accessible and culturable in the lab, at least until the stage at which a potential F0 phenotype can be observed (13). However, it is even more ideal for embryos to be raised to sexual maturity, so that they can be bred to an F2 homozygous null generation. F0 phenotypes, while observable, are known to be more difficult to generate (91). For example, in the apple snail, *P. canaliculata*, there was no observable phenotype in a *pax6* knockout in the F0 KO or F1 heterozygotes, and instead phenotypes were only observed in the F2 generation (17).

There must also be some mode of delivering the guide complex to the embryo, traditionally through microinjection or electroporation (92). In most spiralian KOs, microinjection has been the chosen mode of action but requires training in fine motor skills that can take time to develop (12,13,16,17), and success is likely to be highly species specific. To survive microinjection, embryos must be robust to penetration of the cell membrane by a sharp needle, including the penetration of any additional external membranes such as egg capsules or chorions (93,94). For example, the embryos of the longfin inshore squid, *Doryteuthis pealii*, are surrounded by a protective membrane known as the chorion which microinjection needles cannot pass through (13). To deliver the RNP complex for CRISPR-Cas9 KOs, the chorion must first carefully be cut, and the embryo injected through that hole with bevelled quartz needles (13).

Another important limitation to note is the inability to assess off-target mutations in the genome because of the knock-out (95). In some instances, off-targets can be observed at frequencies higher than the intended mutation, which can have unintended negative effects on the crispant (96). Accurate identification of off-target sites requires access to an annotated genome (95). In the absence of such a resource for *C. fornicata*, CRISPR constructs designed solely from transcriptomic data may lead to unintended edits, and any resulting phenotypes should therefore be interpreted cautiously.

## Conclusions

The production of F0 KO crispant mutants in *C. fornicata* with an observable phenotype and successful genotyping demonstrates that it is possible to use gene-editing techniques in non-traditional model organisms that lack genomic resources, greatly increasing the accessibility of gene-editing to Spiralia and beyond. In the case of the specific cf-*pax6* KO produced here, we propose *C. fornicata* as a genetically enabled embryological system to study the development and evolution of visual systems in molluscs. More broadly, the accessible pipeline presented for transcriptome-driven F0 KO crispant mutant production, will facilitate the study of the genetic control and evolution of diverse physiological traits that are simply absent in many traditional model organisms. Species choice remains a careful decision, as not every non-traditional model organism has robust, readily available embryos that will survive current gene editing delivery techniques (i.e. microinjection or electroporation).

## Data Availability

Raw read data generated in this paper is publicly available at NCBI SRA with the following accession code: PRJNA1333917; raw long read data publicly available at ENA under accession code PRJEB77357; transcriptome assembly and data matrices are available from Biostudies at S-BSST2201; mRNA gene sequence for *cf-pax6* is available from NCBI Genbank, accession number PX425767. File to 3D print microinjection dish mould available at NIH 3D: 3DPX-023156.

## Authors’ Contributions

VAS conceived and supervised all elements of the project. VAS collected samples and extracted RNA for the transcriptome. DCJ and MLR conducted transcriptome assembly. VAS and LB conducted histological analysis of larval and adult eyes. DCJ conducted all CRISPR experiments and analysis. DCJ and VAS drafted the manuscript. SM and TB provided co-supervision of DCJ and reviewed the manuscript. All authors read and approved the final version of the manuscript.

## Acknowledgements

We thank the Marine Biological Laboratory Specimens team for providing adult *Crepidula fornicata* for embryo generation; the Gillis and Rawlinson labs in the Bay Paul Center at the Marine Biological Laboratory for hosting DCJ and VAS while generating CRISPR mutants; Mark Terasaki for his generous support of this work; Grant Batzel for advice on sgRNA and primer design; Stephen Clark at the Babraham Institute for library preparation; Jonathan Henry for microinjection training and providing the 3D printed microinjection dish mould; all members of the Sleight Lab at the University of Aberdeen for technical input and intellectual discussion of this work.

## List of Abbreviations

CRISPR: clustered regularly interspaced palindromic repeats
FSW: filtered sea water
gDNA: genomic DNA
HDR: homology directed repair
ICE: Interference of CRISPR Edits
IDT: Integrated DNA Technologies
KO: knockout
PAM: protospacer adjacent motif
PenStrep: Penicillin G sodium salt /Streptomycin sulphate salt
PBS: phosphate buffer saline
PFA: paraformaldehyde
RNP: ribonucleoprotein
sgRNA: single guide RNA
SNP: single nucleotide polymorphism
TAE: Tris-acetate-EDTA

