## Supplementary Material for "Gene editing without a genome: generation and validation of F0 CRISPR mutants in gastropod mollusc *Crepidula fornicata*"

***Supplemental Table 1.* Percent mutant reads for each embryo from the ICE analysis, with $R^2$ value for model predicting mutations.**

| sgRNA | Single embryo_ID | % mutant reads | $R^2$ value |
|---|---|---|---|
| 17 | 1 | 25 | 0.93 |
| 17 | 2 | 24 | 0.93 |
| 17 | 3 | 15 | 0.95 |
| 17 | 4 | 29 | 0.99 |
| 17 | 5 | 0 | 0.99 |
| 20 | 1 | 46 | 0.91 |
| 20 | 2 | 55 | 0.90 |
| 20 | 3 | 46 | 0.85 |
| 20 | 4 | 67 | 0.90 |

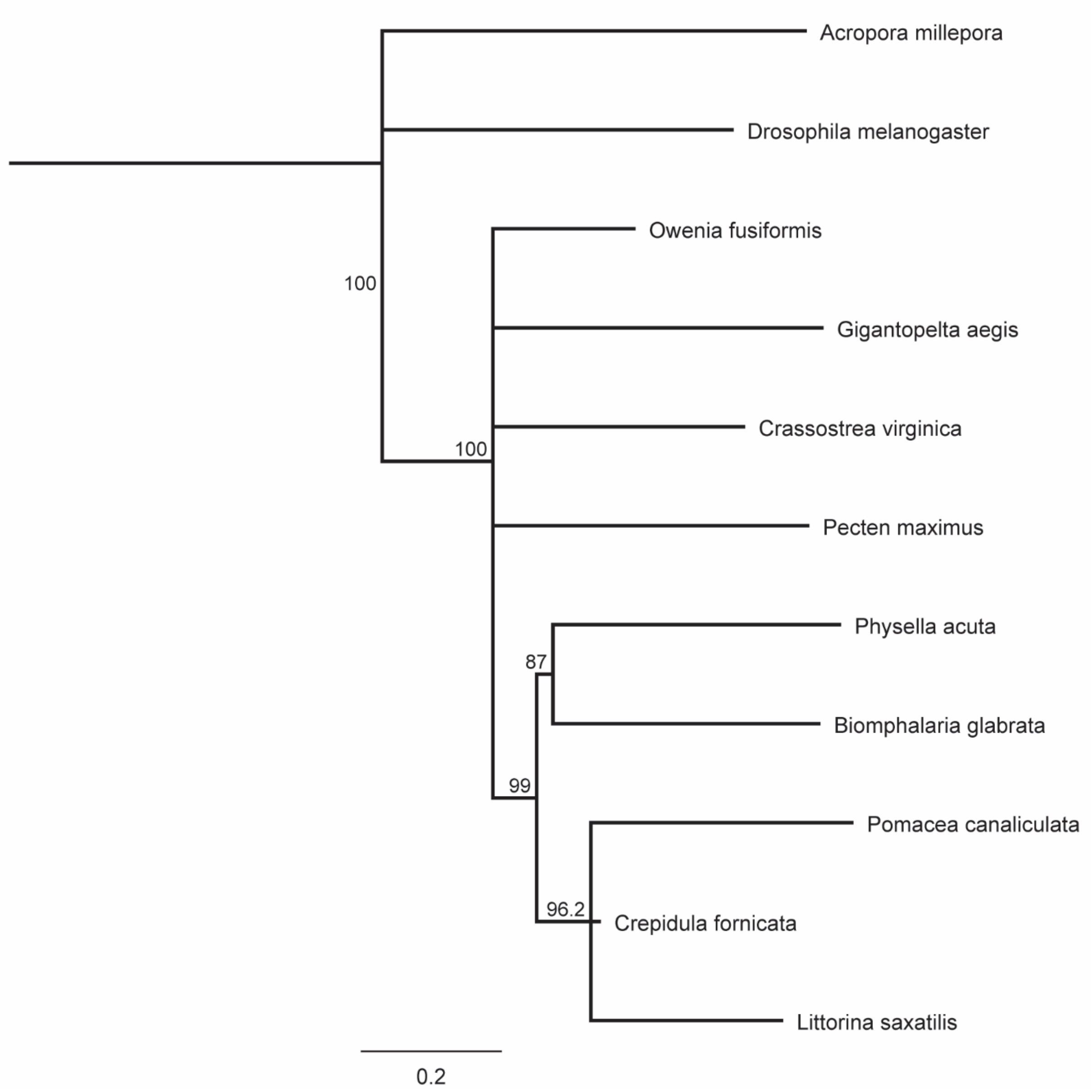


***Supplementary Figure 1*. Phylogenetic relationships between invertebrate *pax6* orthologues.** A neighbour-joining consensus tree showing branch lengths and bootstrap values between *Acropora millepora* (XM_029356362.2), *Drosophila melanogaster* (NM_001272151.1), *Owenia fusiformis* (KY809737.1_cds_AUN27663.1_1), *Gigantopelta aegis* (XM_041510489.1), *Crassostrea virginica* (XM_022437898.1), *Pecten maximus* (XM_033896327.1), *Physella acuta* (XM_059292525.1), *Biomphalaria glabrata* (XM_056044551.1), *Pomacea canaliculata* (XM_025231455.1), *Littorina saxatilis* (XM_070344187.1) orthologues of *pax6*, and the candidate gene, *cf-pax6* (NODE_92639_g45732_i0). Alignments were made using Clustal-Omega, tree was formed in Geneious and exported for visualization.
